# Neurokinin 1 receptor activation in the rat spinal cord maintains latent sensitization, a model of inflammatory and neuropathic chronic pain

**DOI:** 10.1101/2020.05.11.089425

**Authors:** Wenling Chen, Juan Carlos Marvizon

**Affiliations:** Vatche and Tamar Manoukian Division of Digestive Diseases, Department of Medicine, David Geffen School of Medicine at the University of California Los Angeles, Los Angeles, California 90095; Veteran Affairs Greater Los Angeles Healthcare System, Los Angeles, California 90073

**Keywords:** inflammation, nerve injury, neurokinin-1 receptor, primary afferent, substance P

## Abstract

Latent sensitization is a model of chronic pain in which a persistent state of pain hypersensitivity is suppressed by opioid receptors, as evidenced by the ability of opioid antagonists to induce a period of mechanical allodynia. Our objective was to determine if substance P and its neurokinin 1 receptor (NK1R) mediate the maintenance of latent sensitization. Latent sensitization was induced by injecting rats in the hindpaw with complete Freund’s adjuvant (CFA), or by spared nerve injury (SNI). When responses to von Frey filaments returned to baseline (day 28), the rats were injected intrathecally with saline or the NK1R antagonist RP67580, followed 15 min later by intrathecal naltrexone. In both pain models, the saline-injected rats developed allodynia for 2 h after naltrexone, but not the RP67580-injected rats. Saline or RP67580 were injected daily for two more days. Five days later (day 35), naltrexone was injected intrathecally. Again, the saline-injected rats, but not the RP67580-injected rats, developed allodynia in response to naltrexone. To determine if there is sustained activation of NK1Rs during latent sensitization, NK1R internalization was measured in lamina I neurons in rats injected in the paw with saline or CFA, and then injected intrathecally with saline or naltrexone on day 28. The rats injected with CFA had a small amount of NK1R internalization that was significantly higher than in the saline-injected rats. Naltrexone increased NK1R internalization in the CFA-injected rats but nor in the saline-injected rats. Therefore, sustained activation of NK1Rs maintains pain hypersensitivity during latent sensitization.

## 1. Introduction

The latent sensitization model of chronic pain shows that certain injuries put the organism in a long-term state of pain hypersensitivity that is continuously suppressed by opioid and α_2A_ adrenergic receptors (Marvizon et al., 2015; Taylor and Corder, 2014). This shows that chronic pain is a dysregulation of pain pathways so that they stay sensitized after the initial injury is healed. In animals with latent sensitization, opioid receptor antagonists induce episodes of mechanical allodynia and thermal hyperalgesia (Campillo et al., 2011; Corder et al., 2013; Walwyn et al., 2016), demonstrating the coexistence of pain hypersensitivity and its suppression by opioid receptors. Furthermore, mice with genetic deletion of µ-opioid receptors (MORs) do not fully recover from hyperalgesia and no longer respond to MORs antagonists with an episode of allodynia. This occurs both when the MORs deletion is global (Walwyn et al., 2016) or targeted to nociceptive afferents (Severino et al., 2018), indicating that the MORs that suppress the hypersensitivity are located in these afferents.

Different stimuli can induce latent sensitization, including inflammation (Corder et al., 2013; Walwyn et al., 2016), nerve injury (Solway et al., 2011), plantar incision (Campillo et al., 2011; Rivat et al., 2009) or opioids (Rivat et al., 2007; Rivat et al., 2009). However, it is unclear if these represent different types of latent sensitization with different mechanisms.

The initiation, expression and maintenance of latent sensitization are likely to be mediated by different signals: its initiation is probably due to temporary signals evoked by the triggering injury, its maintenance to signals able to endure for long periods of time, and its expression to short-lived signals recruited by the maintenance signals. Differentiating between the maintenance and the expression of latent sensitization is important because drugs that eliminate its maintenance may cure chronic pain. Drugs that inhibit the maintenance of latent sensitization should eliminate the allodynia induced by opioid antagonists permanently, whereas drugs that inhibit its expression should eliminate it only while they are present. In this study, we used this criterion to determine whether neurokinin 1 receptors (NK1Rs) for substance P mediate the expression or the maintenance of latent sensitization.

NK1Rs in dorsal horn neurons are involved in central sensitization (Laird et al., 2001; Xu et al., 1992) and long-term potentiation of the first order synapses of nociceptors (Ikeda et al., 2003; Liu and Sandkuhler, 1998). Substance P release is mediated by NMDA receptors (Marvizon et al., 1997), adenylyl cyclase and protein kinase A (Chen et al., 2018a), which are also required for latent sensitization (Celerier et al., 2000; Corder et al., 2013; Fu et al., 2019). Conversely, substance P release is inhibited by the presynaptic MORs (Chen et al., 2018a) that suppress pain hypersensitivity in latent sensitization (Severino et al., 2018). We hypothesized that NK1Rs mediate the maintenance of latent sensitization and tested this hypothesis by determining whether an NK1R antagonist eliminated latent sensitization. We also determined if NK1Rs are tonically activated by substance P during latent sensitization and if this activation is increased by a MOR antagonist.

## 2. Material and methods

### 2.1 Animals

Male adult (2-4 months old) Sprague-Dawley rats (Envigo, Indianapolis, IN) were used in the study. The Institutional Animal Care and Use Committee of the Veteran Affairs Greater Los Angeles Healthcare System approved all animal procedures, which abide by the Guide for the Care and Use of Laboratory Animals, National Institutes of Health of the United States of America.

### 2.2 Chemicals

Naltrexone (NTX) and RP67580 were from Tocris Bioscience (Minneapolis, MN). Complete Freund’s adjuvant (CFA) and other reagents were from Sigma-Aldrich (St. Louis, MO). RP67580 was dissolved at 10 mM in DMSO and then diluted to 30 µM in saline (final DMSO was 0.3%). NTX was dissolved in saline. CFA was injected undiluted.

### 2.3 CFA injections

Rats were injected subcutaneously in one hindpaw with 50 µl undiluted CFA, using a 26 gauge needle and a 50 µl Hamilton syringe. The needle was inserted in the middle of the paw, near the base of the third toe, at an oblique angle from the heel. It was held for 15 s and then withdrawn.

### 2.4 Spared nerve injury (SNI) by cutting the common peroneal and sural nerves

This is a variation of the SNI model in which the sural nerve is cut instead of the tibial nerve, which decreases the period of mechanical allodynia to less than 30 days (Marvizon et al., 2019; Solway et al., 2011). Rats were anesthetized with isoflurane (2-3 % in O_2_) and prepared for surgery. The skin was cut in the dorsal upper thigh. The muscle was separated to expose the trifurcation of the sciatic nerve. The common peroneal and sural nerves were ligated and cut on both sides of the suture to remove 2-3 mm of each nerve. The muscle was sutured in a continuous pattern. The cut in the skin was closed using a subcuticular intradermal pattern. Rats were given daily injections of the analgesic carprofen for 3 days. Endpoint criteria were motor weakness or signs of paresis, but this did not occur in any of the rats.

### 2.5 Intrathecal (i.th.) injections

Intrathecal catheters were inserted through the lumbar vertebrae (Storkson et al., 1996; Walwyn et al., 2016). Rats were anesthetized with isoflurane (2-3 % in O_2_). Skin and muscle were cut and a 20G needle was inserted between the L5 and L6 vertebrae to puncture the dura mater. The catheter (20 mm of PE-5 tube heat-fused to 150 mm of PE-10 tube) was inserted into the subdural space and pushed rostrally to place its tip at L5-L6, then its PE-10 portion was inserted under the skin of the back and brought out at the head. The skin was sutured. The catheter was filled with saline and heat-sealed. Rats were housed separately. The presence of motor weakness or paresis were criteria for euthanasia, but they did not occur in any of the rats.

Intrathecal injections consisted of 10 µl of injectate followed by 10 μl of saline. Solutions were preloaded into a PE-10 tube and delivered in 1 min. The correct position of the catheter was confirmed postmortem. Three rats lost the catheter after day 28 and were excluded from the measures on day 35.

The dose of NTX, 2.6 nmol i.th., was chosen from our previous studies (Corder et al., 2013; Taylor and Corder, 2014; Walwyn et al., 2016). The dose of the NK1R antagonist RP-67580, 0.3 nmol i.th., was calculated based on its K_i_ of 4.16 nM to displace [^3^H]substance P binding to the NK1R (Garret et al., 1991). The i.th. dose was obtained by multiplying the K_i_ by 30 to get a saturating concentration and then dividing by 500 nM/nmol, the ratio between the EC_50_s for NMDA to induce substance P release in spinal cord slices, 258 nM (Chen et al., 2010), and by i.th. injection *in vivo*, 0.49 nmol (Chen et al., 2014).

### 2.6 Measures of mechanical hyperalgesia

Mechanical allodynia was measured with von Frey filaments (Touch-Test) using the two-out-of-three method (Chen et al., 2018b; Jarahi et al., 2014; Kingery et al., 2000; Michot et al., 2012; Walwyn et al., 2016). Measures were done by putting the rats in acrylic enclosures on an elevated metal grid (IITC Life Science Inc., CA), to which the rats were habituated for 30 min, daily for 3 days. A series of von Frey filaments were applied in ascending order to the plantar surface of the hindpaw for a maximum of 3 s. A withdrawal response was counted if the hindpaw was completely removed from the grid. Each filament was applied three times. The minimal value that caused at least two responses was recorded as the paw withdrawal threshold (PWT). The cut-off threshold was the 15 g filament.

### 2.7 Timeline

The timeline of the experiments to study the effect of the NK1R antagonist RP67580 on latent sensitization is shown in Fig. 1. The rats were habituated to the von Frey enclosures for 3 days. On day -7, the i.th. catheters were implanted. On days -4 and 0, baseline von Frey measures were obtained. On day 0, rats were injected with CFA or underwent SNI. On days 3 and 21, von Frey measures were taken to follow the mechanical allodynia produced by CFA or SNI. On day 28, a baseline von Frey measure was taken, the rats were injected i.th. with vehicle or RP67580, and 15 min later they were injected i.th. with NTX. For the following 2 h, von Frey measures were taken at 15, 30, 60, 120 min after NTX. On days 29 and 30, the rats received daily i.th. injections of vehicle or RP67580. On day 35, a baseline von Frey measure was taken, NTX was injected i.th., and von Frey measures were taken for 2 h at the same time points as before.

**Figure 1.**
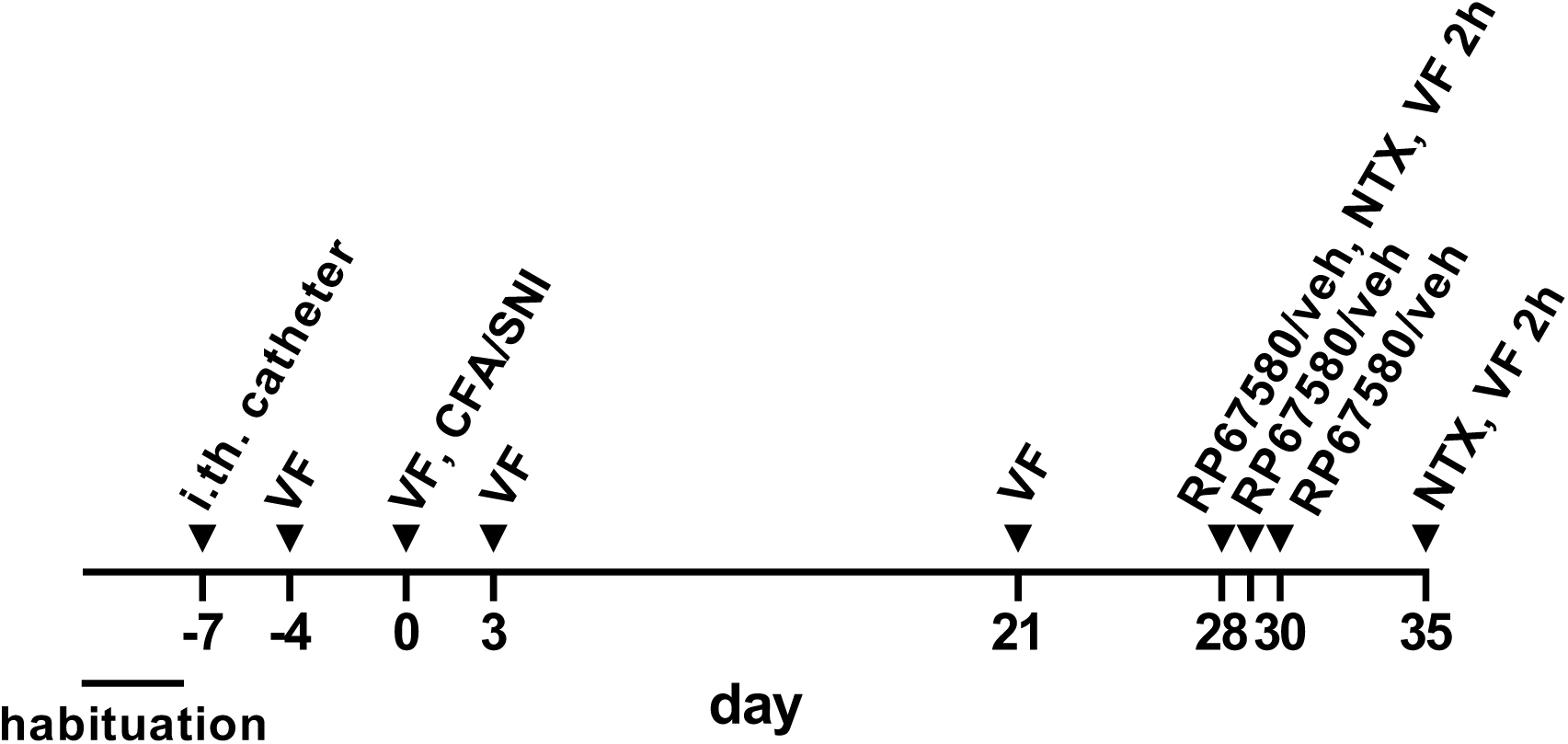
Timeline. Habituation to the von Frey enclosure: 30 min daily for 3 days. Intrathecal (i.th.) catheter implantation. von Frey measures, single (VF) or repeated for 2 h (VF 2h). Complete Freund’s adjuvant (CFA, 50 µl s.c.) injected in the paw. Spared nerve injury (SNI). RP67580, 0.3 nmol i.th. in 10 µl. Vehicle (veh), 10 µl i.th. Naltrexone (NTX) 2.6 nmol i.th. in 10 µl.

### 2.4 Immunohistochemistry

The NK1R antiserum (AB5060, EDM Millipore, Billerica, MA) was raised in rabbit against amino acids 385-407 at the C-terminus of the rat NK1R. This and similar antiserums raised against the same epitope of the NK1R had been fully characterized (Allen et al., 1999; Chen et al., 2018a; Chen et al., 2014; Grady et al., 1996; Honore et al., 1999; Mantyh et al., 1995; Marvizon et al., 2003). They all produce the same staining pattern in the rat spinal cord.

NK1R immunohistochemistry was done as described (Adelson et al., 2009; Marvizon et al., 2003). Rats were killed with pentobarbital (100 mg/kg) and fixed by aortic perfusion of 100 ml 0.1 M sodium phosphate (pH 7.4) containing 0.01% heparin, followed by 400 ml of ice-cold fixative (4% paraformaldehyde, 0.18% picric acid in phosphate buffer). Lumbar spinal cord segments (L4-L5) were post-fixed, cryoprotected, frozen and sectioned at 25 µm using a cryostat. Sections were washed four times and then incubated overnight at room temperature with the NK1R antiserum diluted 1:3000 in phosphate-buffered saline containing 0.3% Triton X-100, 0.001% thimerosal and 10% normal goat serum (Jackson ImmunoResearch Laboratories, West Grove, PA). After three washes, the secondary antibody (1:2000, Alexa Fluor 488 goat anti-rabbit, Invitrogen-Molecular Probes, Eugene, OR) was applied at for 2 h at room temperature. Sections were washed four more times, mounted on glass slides, and coverslipped with Prolong Gold (Invitrogen-Molecular Probes, Eugene, OR).

### 2.5 Quantification of NK1R internalization

NK1R internalization was used as a measure *in situ* of substance P release, a method that has been extensively used and validated (Adelson et al., 2009; Allen et al., 1997; Honore et al., 1999; Mantyh et al., 1995; Marvizon et al., 2003). NK1R neurons in lamina I with or without internalization were visually counted using a Zeiss Axio-Imager A1 (Carl Zeiss, Inc., Thornwood, NY) fluorescence microscope with a 63x (1.40 numerical aperture) objective. The criterion for having internalization was the presence in the neuronal soma of ten or more NK1R endosomes. The person counting the neurons was blinded to the treatment of the rats. Four sections per rat were used, counting all NK1R neurons in lamina I in each section. Results were expressed as the percentage of the NK1R neurons in lamina I with NK1R internalization.

### 2.6 Sample size, randomization and blinding

The reduce the number of animals used, the CFA and SNI control groups were shared with another study on the effect of the Src family kinase inhibitor PP2 on latent sensitization (Chen and Marvizón, 2020). The target sample size was *n* = 7-8 rats per group based on a power analysis. Three rats in the SNI-control group were excluded from measures on day 35 because they had lost the i.th. catheter. Rats were randomly assigned to treatment and drugs. Blinding procedure: solutions of vehicle, PP2 and RP67580 were prepared and randomly coded by JCM. WC injected the coded drugs i.th. and performed the von Frey measures. JCM analyzed the data and un-blinded the results.

### 2.7 Data analysis

Data were analyzed using Prism 8.3 (GraphPad Software, San Diego, CA) and expressed as mean ± standard error of the mean. Statistical significance was set at 0.05. Statistical analyses consisted of repeated-measures two-way ANOVA followed by Holm-Sidak’s post-hoc tests, or three-way ANOVA.

## 3. Results

### 3.1 Effect of the NK1R antagonist on CFA-induced latent sensitization

We studied the effect of the NK1R antagonist RP67580 on the latent sensitization induced by injecting 50 µl CFA in one hindpaw, a model of inflammatory chronic pain. CFA produced strong mechanical allodynia in the ipsilateral hindpaw on day 3, which decreased to near baseline values by days 28-35 (Fig. 2A, D). There was no allodynia in the contralateral hindpaw.

**Figure 2.**
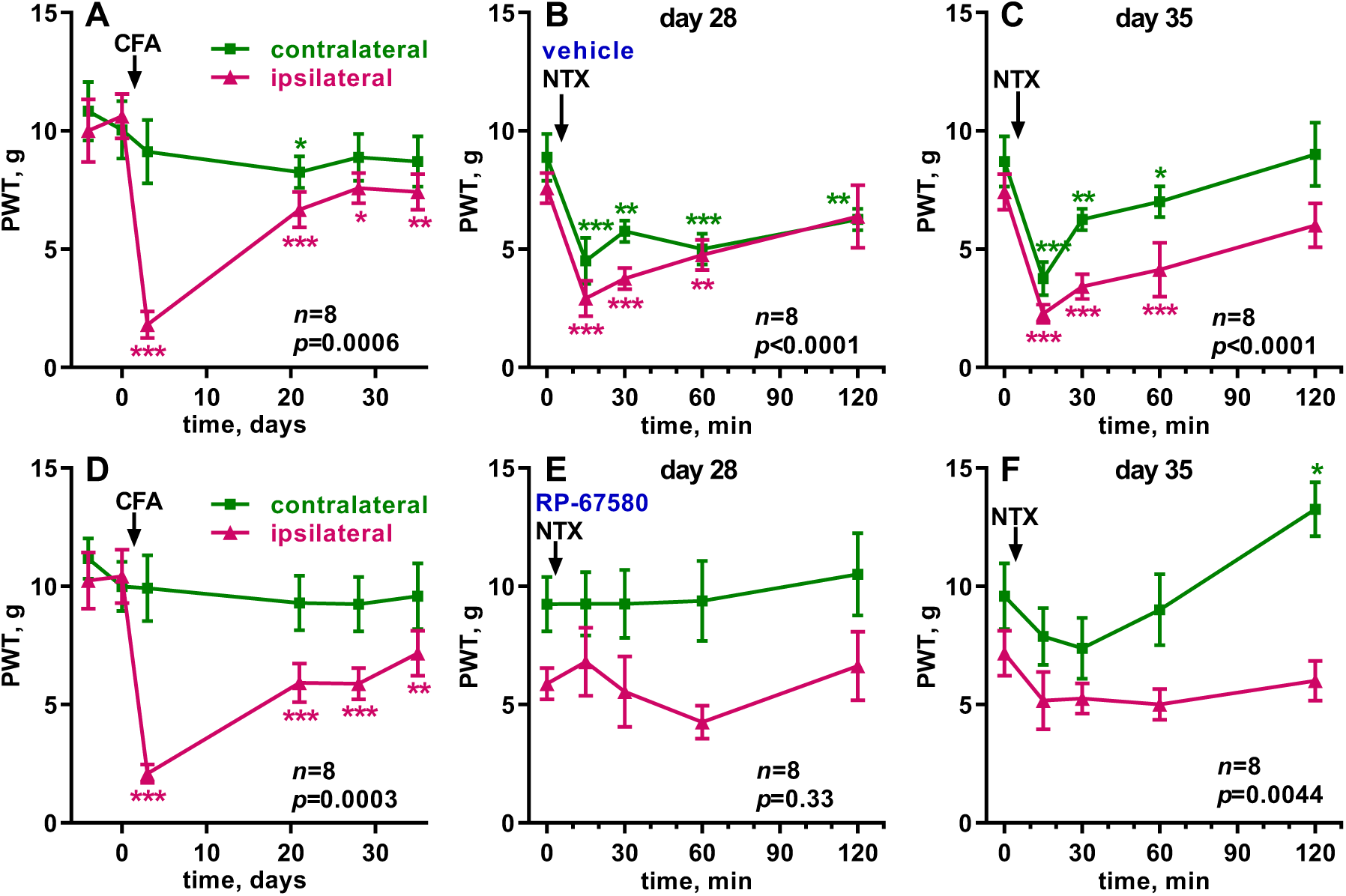
CFA-induced LS, effect of RP67580 on NTX-induced allodynia. Paw withdrawal thresholds (PWT, von Frey) were measured in the ipsilateral (‘ipsi’) and contralateral (‘contra’) hindpaws. **A, D:** CFA (50 µl) was in injected in the hindpaw of eight rats at the time indicated by the arrow. **B, E:** On day 28, 10 µl of vehicle (0.3% DMSO in saline, B) or RP67580 (0.3 nmol in 0.3% DMSO, E) were injected i.th.; NTX (2.6 nmol in 10 µl i.th.) was injected 15 min later (arrow). The following two days, vehicle or RP67580 were injected daily. **C, F:** On day 35, NTX (2.6 nmol in 10 µl i.th.) was injected again (arrow). Each panel was analyzed by RM (both variables) 2-way ANOVA; overall *p* values for ‘time’ (effect of CFA or NTX) are given in each panel. Holm-Sidak’s post-hoc tests comparing to day -4 or 0 min: * *p*<0.05, ** *p*<0.01, *** *p*<0.001.

To study the role of NK1Rs on the expression of latent sensitization, we determined if RP67580 suppressed the allodynia induced by NTX given shortly afterward. Thus, on day 28, rats were injected i.th. with vehicle (control) or RP67580 (0.3 nmol) followed 15 min later by i.th. NTX (2.6 nmol). In the control group, NTX induced bilateral allodynia in the hindpaws, which peaked at 15 min and lasted more than 2 h (Fig. 2B), whereas after RP67580 the NTX-induced allodynia was eliminated (Fig. 2E).

To study the role of NK1Rs on the maintenance of latent sensitization, rats were injected with vehicle or RP67580 daily for two more days and then with NTX (2.6 nmol i.th.) on day 35 (Fig. 1). NTX produced bilateral allodynia in the control rats (Fig. 2C) but not in the rats injected with RP67580 (Fig. 2F). In addition, 2 h after NTX there was an unexpected decrease in responses to the von Frey filaments in the contralateral paw of the rats injected with RP67580.

Results were analyzed by 2-way ANOVA, repeated-measures for the two variables, time and side (Table 1). The variable ‘time’ indicates whether there was a significant effect of CFA or NTX. The variable ‘side’ indicates whether the effects were ipsilateral or bilateral. The effect of CFA over time was significant in both groups (A and D), and a significant effect of ‘side’ and ‘time × side’ showed that the effect of CFA was limited to the ipsilateral paw. On day 28, the effect of NTX over time was significant in the control rats (B) but not in the rats injected with RP67580 (E). On day 35, the effect of NTX over time was significant in the controls (C) and also in the RP67580-injected rats (F), but the later was due to a decrease in allodynia at the 2 h time point and not to the expected increase in allodynia at 15 min and 30 min (Fig. 2F). After the injections of NTX, ‘time × side’ was not significant in the controls, indicating that NTX produced allodynia bilaterally. In the RP67580-injected rats (F), ‘time × side’ was significant on day 35 because the decrease in allodynia at 2 h occurred only in the contralateral paw (Fig. 2F).

**Table 1.**
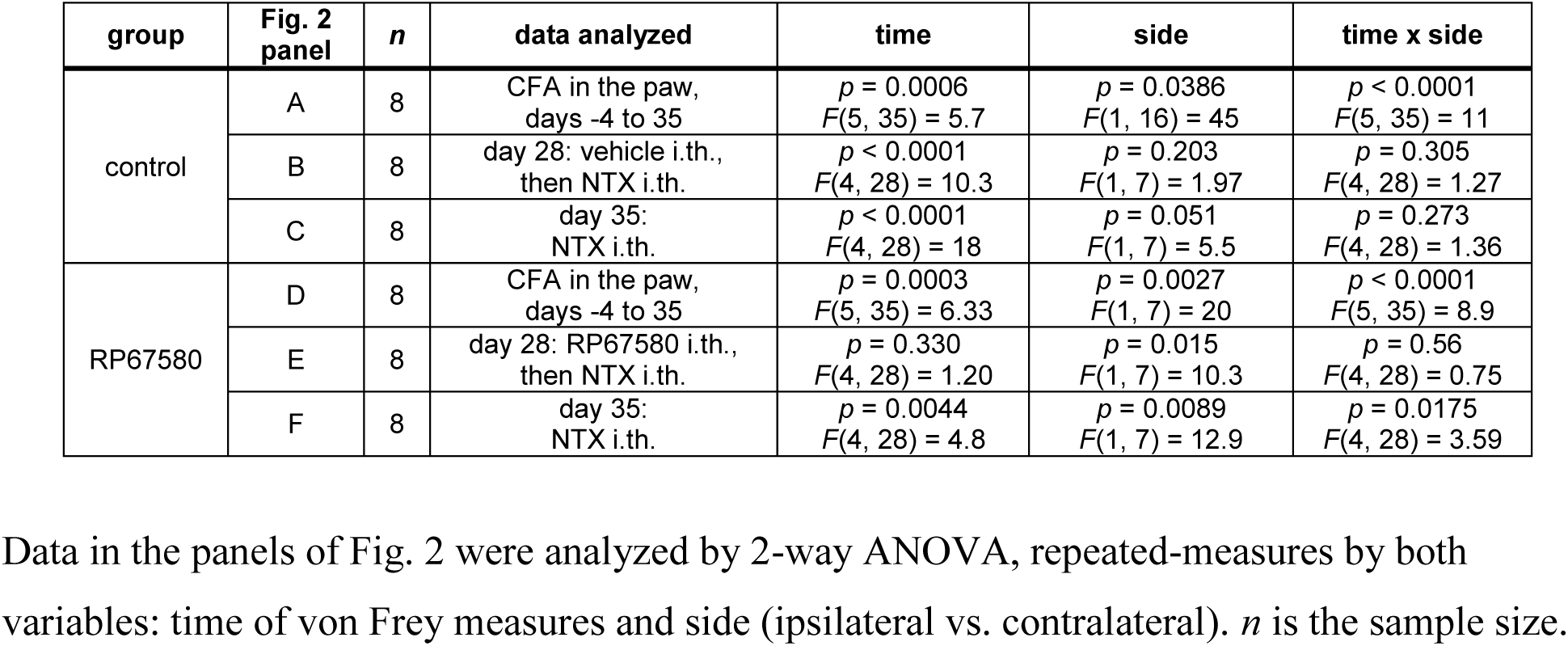
Statistical analyses of data in Fig. 2: CFA-induced latent sensitization.

| group | Fig. 2 panel | n | data analyzed | time | side | time x side |
| --- | --- | --- | --- | --- | --- | --- |
| control | A | 8 | CFA in the paw, days -4 to 35 | $p = 0.0006$<br>$F(5, 35) = 5.7$ | $p = 0.0386$<br>$F(1, 16) = 45$ | $p < 0.0001$<br>$F(5, 35) = 11$ |
| | B | 8 | day 28: vehicle i.th., then NTX i.th. | $p < 0.0001$<br>$F(4, 28) = 10.3$ | $p = 0.203$<br>$F(1, 7) = 1.97$ | $p = 0.305$<br>$F(4, 28) = 1.27$ |
| | C | 8 | day 35: NTX i.th. | $p < 0.0001$<br>$F(4, 28) = 18$ | $p = 0.051$<br>$F(1, 7) = 5.5$ | $p = 0.273$<br>$F(4, 28) = 1.36$ |
| RP67580 | D | 8 | CFA in the paw, days -4 to 35 | $p = 0.0003$<br>$F(5, 35) = 6.33$ | $p = 0.0027$<br>$F(1, 7) = 20$ | $p < 0.0001$<br>$F(5, 35) = 8.9$ |
| | E | 8 | day 28: RP67580 i.th., then NTX i.th. | $p = 0.330$<br>$F(4, 28) = 1.20$ | $p = 0.015$<br>$F(1, 7) = 10.3$ | $p = 0.56$<br>$F(4, 28) = 0.75$ |
| | F | 8 | day 35: NTX i.th. | $p = 0.0044$<br>$F(4, 28) = 4.8$ | $p = 0.0089$<br>$F(1, 7) = 12.9$ | $p = 0.0175$<br>$F(4, 28) = 3.59$ |

### 3.2 Effect of the NK1R antagonist on SNI-induced latent sensitization

We also studied the effect of the NK1R antagonist on the latent sensitization induced by SNI, a model of neuropathic pain. To get a faster return to baseline responses, SNI was induced by cutting the common peroneal and sural nerves (Solway et al., 2011) instead of the most common procedure of cutting the common peroneal and tibial nerves (Marvizon et al., 2019). SNI produced strong mechanical allodynia in the ipsilateral hindpaw on day 3, which went back to baseline values by day 28 (Fig. 3A, D). There was no allodynia in the contralateral hindpaw.

**Figure 3.**
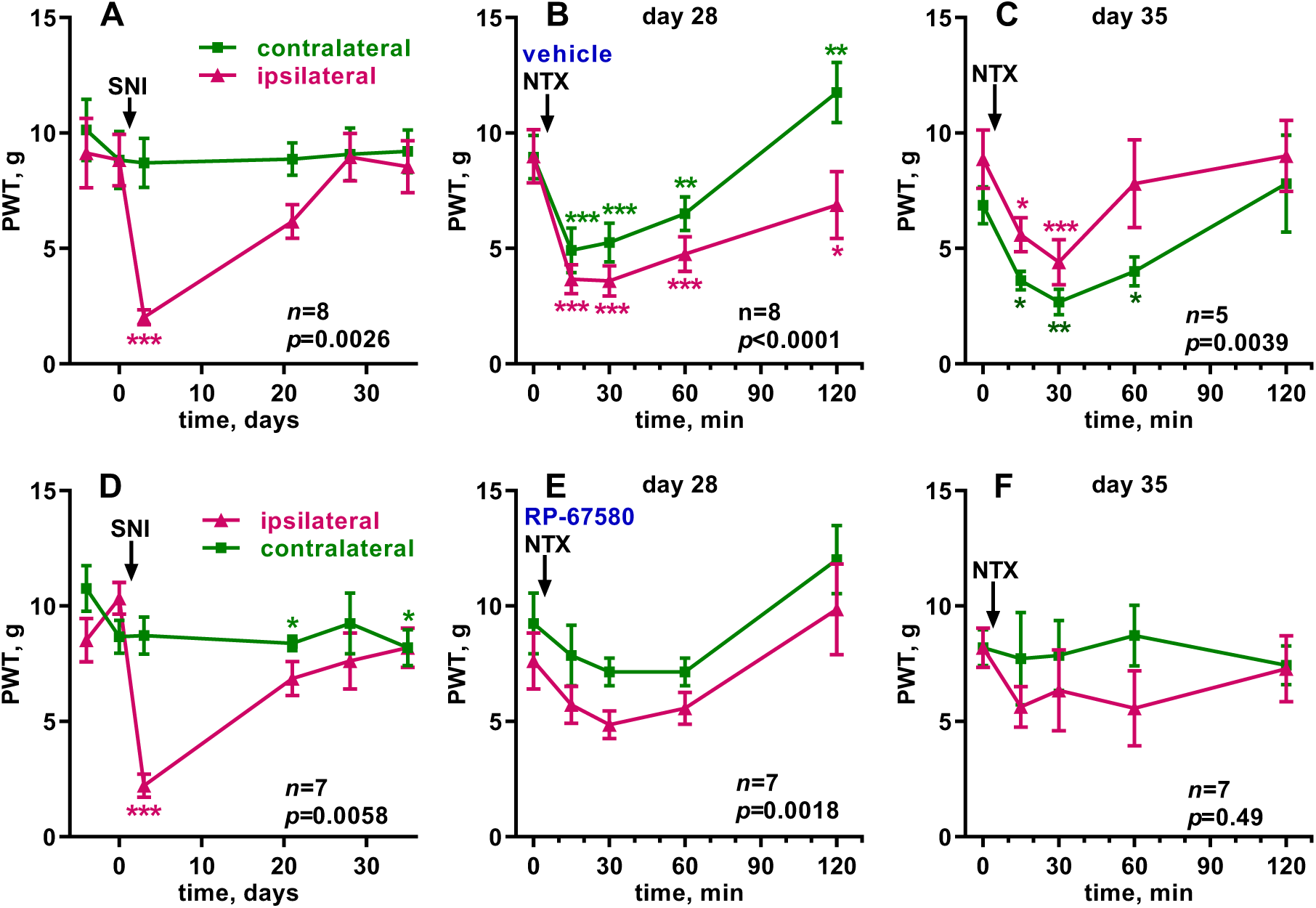
SNI-induced LS, effect of RP67580 on NTX-induced allodynia. Paw withdrawal thresholds (PWT, von Frey) were measured in the ipsilateral (‘ipsi’) and contralateral (‘contra’) hindpaws. **A, D:** SNI was performed in one hindpaw of eight (A-C) or seven (D-F) rats at the time indicated by the arrow. **B, E:** On day 28, 10 µl of vehicle (0.3% DMSO in saline, B) or RP67580 (0.3 nmol in 0.3% DMSO, E) were injected i.th.; NTX (2.6 nmol in 10 µl i.th.) was injected 15 min later (arrow). The following two days, vehicle or RP67580 were injected daily. **C, F:** On day 35, NTX (2.6 nmol in 10 µl i.th.) was injected again (arrow). Each panel was analyzed by RM (both variables) 2-way ANOVA; overall *p* values for ‘time’ (effect of CFA or NTX) are given in each panel. Holm-Sidak’s post-hoc tests comparing to day -4 or 0 min: * *p*<0.05, ** *p*<0.01, *** *p*<0.001.

To study the role of NK1Rs on the expression of SNI-induced latent sensitization we determined if RP67580 suppressed the allodynia induced by NTX given shortly afterward. On day 28, rats were injected i.th. with vehicle (control) or RP67580 (0.3 nmol), and 15 min later with NTX (2.6 nmol). In the control group, NTX induced bilateral allodynia in the hindpaws, which peaked at 15 min and lasted 2 h (Fig. 3B). After RP67580, the NTX-induced allodynia was largely eliminated (Fig. 3E).

To study the role of NK1Rs on the maintenance of SNI-induced latent sensitization, rats were injected i.th. with vehicle or RP67580 daily for two more days and then with NTX (2.6 nmol i.th.) on day 35 (Fig. 1). NTX produced bilateral allodynia in the control rats (Fig. 3C) but not in the rats injected with RP67580 (Fig. 3F).

Results were analyzed by repeated-measures 2-way ANOVA (Table 2). The effect of SNI over time was significant in both groups (A and D), and a significant effect of ‘side’ and ‘time × side’ showed that the effect of SNI was limited to the ipsilateral paw. On day 28, the effect of NTX over time was significant in the control rats (B) and also in the RP67580-injected rats (E), but the later seems to be due both to a small increase in allodynia at 30 min and to a decrease in allodynia at 2 h, both of which were not statistically significant in the Holm-Sidak’s post-hoc test (Fig. 3E). On day 35, the effect of NTX over time was significant in the controls (C) but not in the rats injected with RP67580 (F). After the injections of NTX, ‘time × side’ was not significant in the RP67580-injected rats (E, F), indicating a lack of effect of NTX in either paw. In the control rats, the effect of ‘time × side’ was significant on day 28 due to a decrease in allodynia in the contralateral paw at 2 h (Fig. 3B). The effect of ‘time × side’ was not significant in the control rats on day 35 (Fig. 3C).

**Table 2.** Statistical analyses of data in Fig. 3: SNI-induced latent sensitization.

| group | Fig. 3 panel | n | data analyzed | time | side | time x side |
| --- | --- | --- | --- | --- | --- | --- |
| control | A | 8 | SNI, days -4 to 35 | $p = 0.0026$<br>$F(5, 35) = 4.58$ | $p = 0.0017$<br>$F(1, 7) = 24$ | $p < 0.0001$<br>$F(5, 29) = 8.3$ |
| | B | 8 | day 28: vehicle i.th., then NTX i.th. | $p < 0.0001$<br>$F(4, 28) = 16$ | $p = 0.0085$<br>$F(1, 7) = 13$ | $p = 0.0042$<br>$F(4, 28) = 4.8$ |
| | C | 5 | day 35: NTX i.th. | $p = 0.0039$<br>$F(4, 16) = 6.0$ | $p = 0.063$<br>$F(1, 4) = 27$ | $p = 0.403$<br>$F(4, 16) = 1.07$ |
| RP67580 | D | 7 | SNI, days -4 to 35 | $p = 0.0058$<br>$F(5, 30) = 4.12$ | $p = 0.0006$<br>$F(1, 6) = 42$ | $p < 0.0001$<br>$F(5, 30) = 9.4$ |
| | E | 7 | day 28: RP67580 i.th., then NTX i.th. | $p = 0.0018$<br>$F(4, 24) = 5.9$ | $p = 0.0084$<br>$F(1, 6) = 14.9$ | $p = 0.98$<br>$F(4, 24) = 0.08$ |
| | F | 7 | day 35: NTX i.th. | $p = 0.49$<br>$F(4, 24) = 0.88$ | $p = 0.14$<br>$F(1, 6) = 2.88$ | $p = 0.52$<br>$F(4, 24) = 0.82$ |

### 3.3 Substance P release and NK1R activation during latent sensitization

If latent sensitization is maintained by NK1Rs in dorsal horn neurons, then these receptors should be activated by a sustained release of their agonists substance P and neurokinin A. This should result in increased basal NK1R internalization in lamina I neurons in rats with latent sensitization. Substance P release from primary afferents is inhibited by MORs (Chen et al., 2018a; Kondo et al., 2005), which are constitutively active during latent sensitization (Corder et al., 2013; Walwyn et al., 2016). Therefore, we predicted that an i.th. injection of NTX to rats with latent sensitization would increase NK1R internalization.

To test these two predictions, we injected rats in the hindpaw with 50 µl CFA to induce latent sensitization. Control rats were injected with 50 µl saline. The control rats developed a small amount of bilateral mechanical allodynia on day 7 (Fig. 4A), which resulted in a significant effect of the variable time (Table 3A). In contrast, the CFA-injected rats showed strong allodynia ipsilaterally on days 3-7, which gradually returned to baseline levels (Fig. 4B). No allodynia was observed in the contralateral hindpaw. The CFA-injected rats showed a significant effect of CFA over time, significant differences in the response of the hindpaws (‘side’) and a significant interaction of these two variables (Table 3B). On day 28, the control and CFA-injected groups were divided into two sub-groups: one that received i.th. saline and another that received i.th. NTX (2.6 nmol). Rats were killed and fixed 30 min after the i.th. injection, and lumbar (L4-L5) segments of the spinal cord were processed for NK1R immunohistochemistry.

**Table 3.**
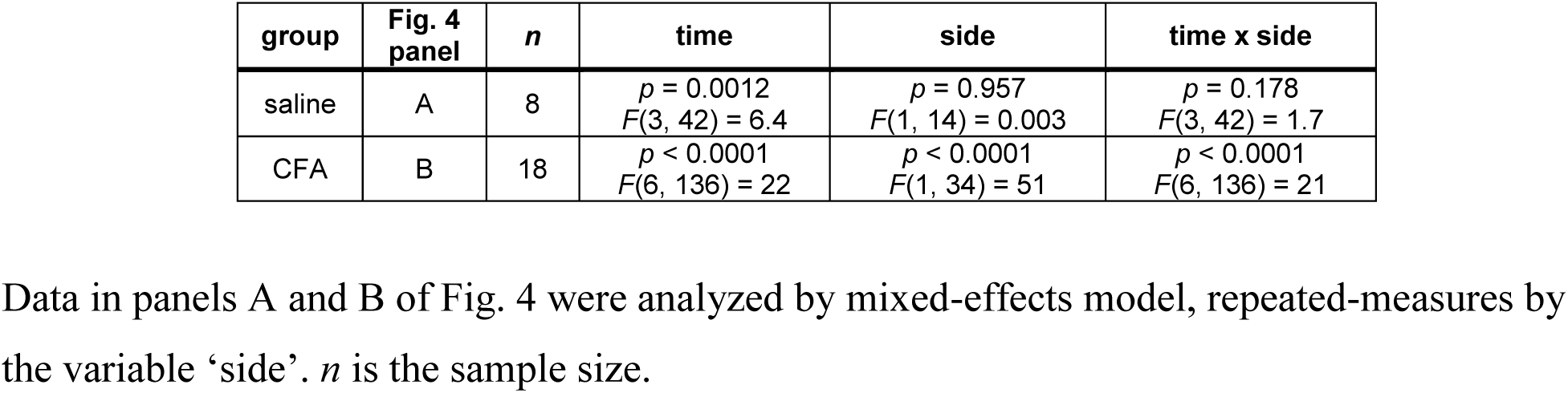
Statistical analyses of von Frey data in Fig. 4.

**Figure 4.**
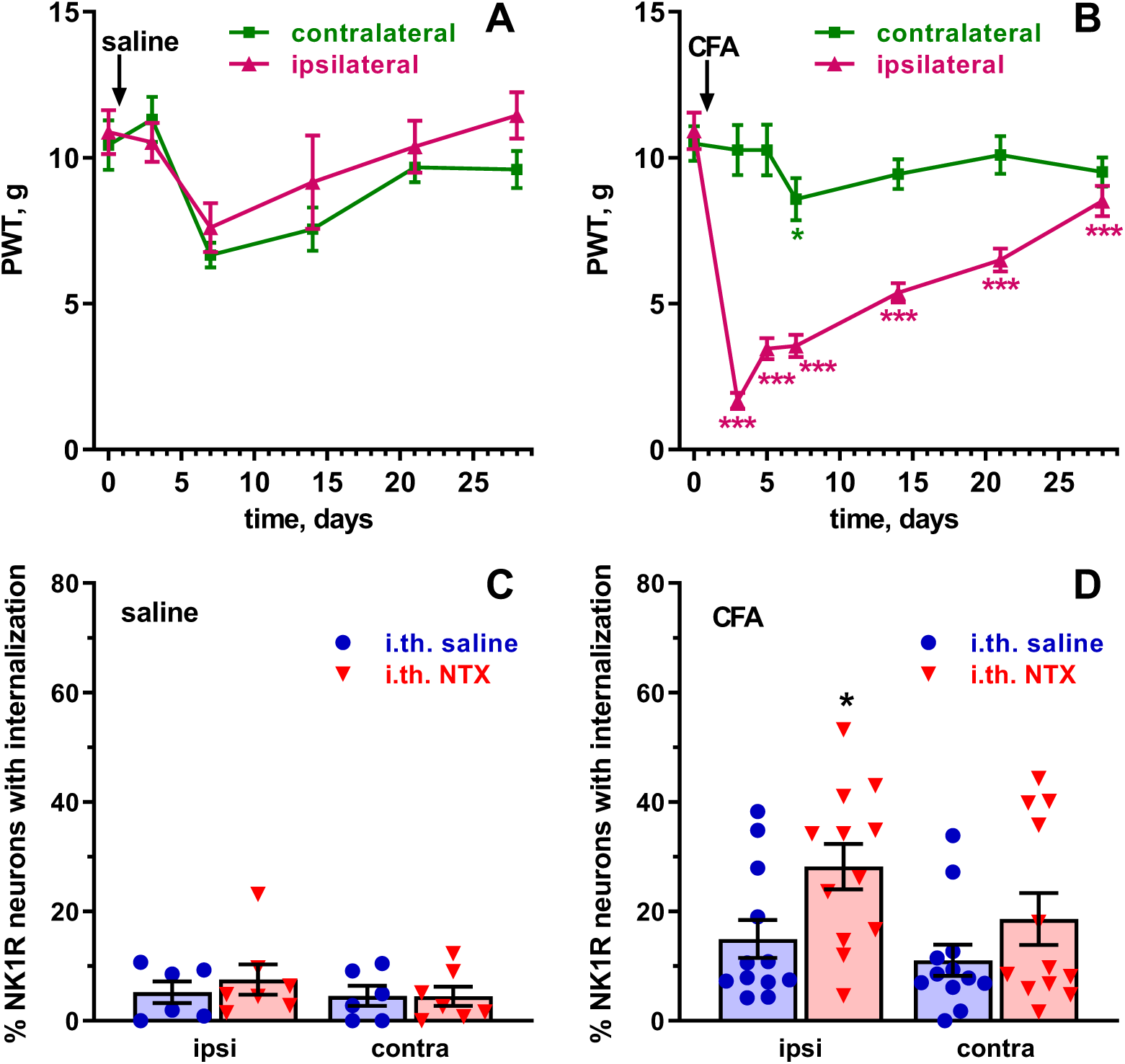
NK1R internalization during latent sensitization. Rats were injected in one hindpaw with 50 µl saline (**A**, n=14 rats) or CFA (**B**, n=24 rats). Paw withdrawal thresholds (PWT, von Frey filaments) were measured in the ipsilateral and contralateral hindpaws at the times indicated. On day 28, rats injected in the paw with saline (**C**, *n*=6 or 7 rats/group) or CFA (**D**, *n*=12 rats/group) received 10 µl intrathecal saline or NTX (2.6 nmol). Rats were killed 30 min later and NK1R internalization was measured in lamina I neurons of the L4-L5 dorsal horns ipsilateral and contralateral to the injected paw. Analyses are shown in Tables 3 and 4. Holm-Sidak’s post-hoc tests comparing to time 0 (A, B) or i.th. saline (C, D): * *p*<0.05, *** *p*<0.001.

Negligible NK1R internalization was found in the control rats without latent sensitization, regardless of whether they received i.th. saline or i.th. NTX (Fig. 4C). In contrast, rats with CFA-induced latent sensitization showed significant amounts of NK1R internalization even after i.th. saline, and i.th. NTX increased the number of neurons with NK1R internalization (Fig. 4D). A three-way ANOVA of the combined data of panels C and D of Fig. 4 (Table 4) yielded significant effects for the variables CFA and NTX, showing that both the CFA injection in the paw and i.th. NTX resulted in significant increases in NK1R internalization.

**Table 4.**
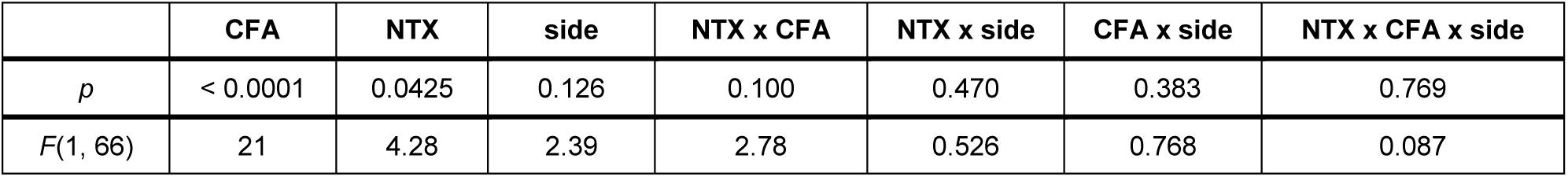
Three-way ANOVA of NK1R internalization data in Fig. 4C, D.

In previous work we have provided numerous confocal images of NK1R neurons in the dorsal horn with and without NK1R internalization (Adelson et al., 2009; Chen et al., 2014; Marvizon et al., 1997; Zhang et al., 2010). The NK1R neurons examined in this study were entirely similar to the ones shown in those reports.

These results confirmed our two predictions and strongly support the ideas that 1) there is sustained substance P release during latent sensitization and 2) this substance P release is partially, but not completely, suppressed by MORs.

## 4. Discussion

### 4.1 NK1Rs mediate the maintenance of latent sensitization

We postulate that different signals mediate the initiation, the maintenance and the expression of latent sensitization. In it, hyperalgesia is suppressed by opioid receptors (Corder et al., 2013; Severino et al., 2018; Walwyn et al., 2016). Therefore, if NTX-induced allodynia is eliminated several days after injecting a compound, then it must have eliminated latent sensitization by blocking a maintenance mechanism. In contrast, inhibitors of mechanisms that mediate the expression of latent sensitization would inhibit NTX-induced allodynia only while the compound is present.

We found that the NK1R antagonist RP67580 eliminated the allodynia induced by NTX not only when it was given shortly afterward, but also when NTX was given one week later. We used three daily i.th. injections of SR67580 and did not assess if a single injection of SR67580 would achieve the same effect. Also, it is possible that NTX-induced allodynia returns after one week. Still, our findings provide a strong indication that NK1Rs are involved in the maintenance of latent sensitization. In a parallel study, we used the same criterion to show that a Src family kinase also maintains latent sensitization (Chen and Marvizón, 2020).

Our results are consistent with a previous study (Rivat et al., 2009) showing that selective elimination of NK1R-expressing neurons in the dorsal horn with substance P-saporin decreased the mechanical allodynia and thermal hyperalgesia induced by paw incision and several injections of the opioid fentanyl, two stimuli that induce latent sensitization (Campillo et al., 2011; Lian et al., 2010; Rivat et al., 2007). However, Rivat et al. (2009) studied only the initial period of hyperalgesia induced by incision or fentanyl (up to day 11) and not the reinstatement of hyperalgesia by NTX, the hallmark of latent sensitization.

### 4.2 Mechanism

Neurons with NK1Rs in lamina I of the dorsal horn are key components of the dorsal horn circuitry that modulates pain in rodents (Gutierrez-Mecinas et al., 2016; Haring et al., 2018; Polgar et al., 2013; Sathyamurthy et al., 2018; Todd, 2017; Yasaka et al., 2014) because many of them send axons to the parabrachial nucleus and the brainstem (Polgár et al., 2010; Ruscheweyh et al., 2004; Todd et al., 2000). These neurons receive direct synapses from peptidergic C/Aδ fibers (Todd et al., 2000; Torsney and MacDermott, 2006), and these synapses are an important locus of pain regulation (Li et al., 2018). NK1R neurons also receive synapses from vertical cells, which serve as a hub to integrate signals from excitatory and inhibitory interneurons of the dorsal horn (Kato et al., 2009; Peirs and Seal, 2016; Peirs et al., 2015; Yasaka et al., 2010; Yasaka et al., 2014). Activation of NK1Rs in dorsal horn neurons produces slow depolarizations, increases excitability (Murase and Randic, 1984; Murase et al., 1986), increases NMDA receptor currents (Rusin et al., 1993; Rusin et al., 1992), and induces long-term potentiation of their synapses with primary afferents (Ikeda et al., 2003; Liu and Sandkuhler, 1997; Liu and Sandkuhler, 1998). Therefore, sensitization of NK1R neurons in lamina I would increase the gain of pain signals sent to the brain, which can explain the pain hypersensitivity during latent sensitization.

### 4.3 NK1Rs are involved in both inflammatory and neuropathic pain

The NK1R antagonist RP67580 eliminated latent sensitization induced by either CFA or SNI, indicating that NK1Rs maintain latent sensitization induced by inflammation or nerve injury. Previous studies have shown that NK1R expression in dorsal horn neurons increases after inflammation induced by CFA, arthritis and nerve injury (Abbadie et al., 1997; Allen et al., 1999; Honore et al., 1999; Krause et al., 1995). Cahill and Coderre (2002) showed that the NK1R antagonist L-732,138, given i.th. daily for 4 days, inhibited the induction of mechanical and cold allodynia produced by sciatic nerve constriction. L-732,138 also decreased allodynia when given at days 8-12 after nerve injury, but its effect was gone by day 16. Hence, NK1Rs do not seem to maintain allodynia induced by nerve constriction, at least at these early time points. The initial phase of allodynia induced by nerve injury could be mediated by mechanisms different from those that maintain latent sensitization.

### 4.4 Sustained substance P release and NK1R activation during latent sensitization

If latent sensitization is maintained by the continuous activation of NK1Rs, this should be driven by a sustained release of substance P, since there is no evidence for NK1R constitutive activity. Indeed, we found that NK1R internalization is higher in rats with latent sensitization than in control rats, indicating that there is increased activation of NK1Rs during latent sensitization. However, the amount of NK1R internalization was quite low, ∼15%, showing that a few active NK1Rs are sufficient to maintain hypersensitivity. This suggests that a complete blockade of NK1Rs by an antagonist is necessary to eliminate latent sensitization. NTX increased the number of neurons with NK1R internalization to ∼30% in the CFA-injected rats, consistent with the idea that MORs in primary afferents suppress hyperalgesia during latent sensitization (Severino et al., 2018; Walwyn et al., 2016) by inhibiting the release of substance P and other neurotransmitters.

These findings are consistent with previous studies. Thus, NK1Rs are tonically active during the second phase of the formalin test (Henry et al., 1999). CFA injection in the paw or the joint, and nerve injury, increases the expression of NK1Rs and their internalization after electrical stimulation of primary afferents (Abbadie et al., 1997; Allen et al., 1999; Honore et al., 1999).

### 4.5 Failure of NK1R antagonists in clinical trials

Any report on the anti-hyperalgesic effect of NK1R antagonists inevitably elicits the criticism that these antagonists have failed as analgesics in clinical trials. Nevertheless, the evidence presented in this study and previous work clearly shows that NK1Rs in dorsal horn neurons play a key role in pain modulation in rodents. This discrepancy was examined in two brief reviews 20 years ago (Hill, 2000; Urban and Fox, 2000), but no further clinical trials on NK1R antagonists as analgesics were performed.

There may be species differences in the role of NK1Rs between rodents and humans. NK1R immunoreactive neurons in the human spinal cord are similar to the ones in the rat (Ding et al., 1999), so it is likely that dorsal horn neurons with NK1Rs are also projection neurons in humans. Still, it is possible that in humans the spinothalamic track is more relevant for pain perception than the pathways connecting the spinal cord to the brainstem and the parabrachial nucleus originating in the NK1R neurons (Todd et al., 2000). If so, the critical differences in pain pathways between rodents and humans need to be studied.

Hill (2000) suggested that the analgesic effects of NK1R antagonists could be due to their supraspinal effects on stress (Rupniak and Kramer, 2017). Indeed, stress induces hyperalgesia in rats with latent sensitization (Chen et al., 2018b; Rivat et al., 2007). However, in this study both RP67580 and NTX were given intrathecally, so it is unlikely that they had supraspinal effects. This volume of intrathecal injection precludes the injectate from reaching the brain (Yaksh and Rudy, 1976).

Another possibility is that the clinical trials with NK1R antagonists may not have detected the effects on chronic pain studied here. One of these clinical trials (Dionne et al., 1998) studied the effect of the NK1R antagonist CP-99,994 on pain measured up to 4 h after a dental extraction, which is not a measure of chronic pain. Nevertheless, CP-99,994 produced significant analgesia at 90 min. Three other clinical trials studied the effect of the NK1R antagonist lanepitant. The effect of lanepitant on migraine (Goldstein et al., 2001a) was close to statistical significance (*p*=0.065). Lanepitant did not decrease pain in osteoarthritis (Goldstein et al., 2000). Osteoarthritis is a chronic pain condition but, unlike latent sensitization, it involves a persistent peripheral injury. Lanepitant also failed to alleviate diabetic neuropathy (Goldstein et al., 2001b), perhaps because it is mechanistically different from nerve transection. In all three studies, lanepitant was administered orally and it is unclear whether it can cross the blood-brain barrier. NK1R antagonists have important effects in the brain (Rupniak and Kramer, 2017) that might counteract their effects on the spinal cord. Finally, the discovery of biased agonism (Simmons, 2005; Smith et al., 2018; Urban et al., 2007) shows that the properties of one agonist cannot be extended to other agonists at the same receptor.

The latent sensitization model shows that the chronic pain state is radically different from normal pain transmission. The goal, then, is not so much to reduce pain but to reset the organism to the normal state. This study indicates that NK1R antagonists may be able to do this. Our strategy may be useful to design any future clinical trials.

### 4.6 Conclusions

Activation of NK1Rs in lamina I neurons of the dorsal horn maintains pain hypersensitivity in the latent sensitization model of chronic pain. This is true for latent sensitization initiated by either inflammation or nerve injury.

## Abbreviations

ANOVA: analysis of variance
CFA: complete Freund’s adjuvant
EC_50_: dose that produces half the effect
i.th.: intrathecal
K_i_: equilibrium inhibition constant
MOR: µ opioid receptor
NTX: naltrexone
NK1R: neurokinin 1 receptor
NMDA: N-methyl-D-aspartate
PE: polyethylene
PWT: paw withdrawal threshold
RP67580: (3aR,7aR)-Octahydro-2-[1-imino-2-(2-methoxyphenyl)ethyl]-7,7-diphenyl-4H-isoindol
SNI: spared nerve injury

